# Mitogenome reconstruction of an endangered African seahorse from a Traditional Chinese Medicine market was based on a misidentification

**DOI:** 10.1101/2020.07.14.202978

**Authors:** Peter R. Teske

## Abstract

The recently published complete mitochondrial genome of the endangered Knysna seahorse, *Hippocampus capensis* Boulenger, 1900, was based on a specimen obtained from a Traditional Chinese Medicine market. As *H. capensis* is endemic to temperate South Africa and exceptionally rare, illegal trade to supply Asian markets would constitute a considerable extinction risk. I investigated the phylogenetic placement of the Chinese specimen using mitochondrial DNA control region and cytochrome *b* sequences from the *H. capensis* mitogenome among corresponding published sequences of *H. capensis* and a number of closely related seahorse species. The Chinese specimen was distinct from *H. capensis* and instead clustered with *H. casscsio*, a seahorse that was recently described from the South China Sea. The sequences of *H. casscsio* clustered randomly among those of specimens identified as *H. fuscus*, a species whose taxonomic validity is disputed, and which is considered to be a synonym of the widespread Indo-Pacific seahorse *H. kuda*. Given that the morphological identification of seahorses is difficult, it is recommended that the publication of new species descriptions and genomic resources be preceded by a comprehensive comparison with the available molecular data. The taxonomy of seahorses is far from resolved, and cutting-edge molecular studies will not improve this situation if they do not take existing information into consideration.

## Introduction

Seahorses (genus *Hippocampus* Rafinesque, 1810) are teleost fishes from the family Syngnathidae which have a highly specialized morphology that includes an upright posture, a prehensile tail, fused jaws, an exoskeleton and a male brood pouch. While these unique morphological features make seahorses easy to recognize, distinguishing between different species is difficult because of considerable morphological variation that is affected by factors such as age, gender and the environment (Lourie et al. 2016). As a result, a large number of synonyms exist, and the number of species accepted as valid in recent taxonomic treatments have ranged from 23 to 83 species (Lourie et al. 2016).

Molecular methods represent a useful independent means of confirming species status that requires little taxonomic expertise (Hebert et al. 2003), and such methods are now often used to support the distinctness of newly described species from morphologically similar seahorses (González et al. 2014; Zhang et al. 2016; Short et al. 2020), or challenge the validity of new species descriptions (Teske et al. 2007). While most studies have used the DNA barcoding marker for animals, the mitochondrial DNA cytochrome *c* oxidase subunit I gene (COI or cox1), advances in DNA sequencing technology have now made it possible to sequence the entire mitochondrial genome (or mitogenome), and several seahorse mitogenomes have recently been published (Zhang et al. 2015; Wang et al. 2016, 2019b; Chen et al. 2018; Ge et al. 2018; Lai et al. 2019; Jahari et al. 2020). The increased level of resolution provided by mitogenomes can be expected to significantly improve taxonomic resolution in cases where DNA barcoding provides insufficient information for distinguishing closely related species.

One of the recently published seahorse mitogenomes, that of the endangered Knysna seahorse, *Hippocampus capensis* Boulenger, 1900 (Ge et al. 2018), was reconstructed using DNA of a specimen obtained from the Bozhou Chinese herbal medicine market (Bozhou, Anhui Province, PR China). The import of African seahorses for Traditional Chinese Medicine (TCM) markets in Asia is very common, and has resulted in a serious depletion of local stocks (T. Mkare, KMFRI, Mombasa, Kenya and I. da Silva, Universidade Lúrio, Pemba, Mozambique, pers. comm.). For example, the mitogenome of the East African *H. camelopardalis* was generated using a specimen from the same TCM market in Bozhou (Lai et al. 2019), and both the West African seahorse *H. algiricus* and the South Africa *H. capensis* were reported from a Taiwanese TCM market (Chang et al. 2013). The sale of *H. capensis* at these markets is a concern because this species is endangered, and endemic to three estuaries on the temperate south coast of South Africa (Teske et al. 2003; Mkare et al. 2017). The illegal collection of these seahorses for TCM markets can be expected to rapidly drive this exceptionally rare species to extinction.

The morphological identification of the Chinese specimen was relatively vague, and based on the absence of a coronet, a short snout and the absence of spines on the body. A comparison of the image provided in the mitogenome announcement with *H. capensis* suggested a questionable identification, with the difference in snout length being the most obvious morphological discrepancy (Teske et al. 2005). In contrast, the specimen of *H. capensis* from a TCM in Taiwan (Chang et al. 2013) resembles South African *H. capensis* more closely. To confirm the suspected distinctness of the Chinese seahorse used for the mitogenome reconstruction from the South African *H. capensis*, I compared its DNA sequences of partial mitochondrial control region and cytochrome *b* sequences with those of *H. capensis* and a number of closely related species. The reason for selecting these two markers is that they are available from a comparatively large number of species and locations.

## Methods

Sequences were downloaded from GenBank and aligned in MEGA7 (Kumar et al. 2016) using the ClustalW alignment tool (Thompson et al. 2002). Consensus sequences 402 bp (control region) and 836 bp (cytochrome *b*) in length that were present in most sequences were then used to reconstruct phylogenetic trees using the neighbour-joining algorithm (Saitou and Nei 1987) in MEGA7. In both cases, the Tamura 3-parameter model (Tamura 1992) with gamma distribution was identified as the most suitable model of nucleotide evolution using the Bayesian Information Criterion (Schwarz 1978). Pairwise deletion of missing data was applied, and nodal support was based on 1000 bootstrap replications (Felsenstein 1985).

## Results and Discussion

Both phylogenetic trees recovered previously published *Hippocampus capensis* sequences (some of which I collected myself and can thus vouch for their authenticity) as well supported monophyletic clusters. The Chinese specimen used for the mitogenome reconstruction of *H. capensis* was not part of these clusters, but was grouped with sequences from the recently described *H. casscsio* Zhang, Qi, Wang & Lin, 2016, specimens identified as *H. fuscus* Rüppell, 1838 from the Mediterranean, the Red Sea and, in the case of cytochrome *b* data, the Arabian Sea, as well as *H. kuda* Bleeker, 1852.

These results confirm that the Chinese specimen was misidentified. The portion of the mitochondrial control region of this specimen is identical to one of the sequences of *H. casscsio*, from Beibu Bay, South China Sea (Zhang et al. 2016), whereas its partial cytochrome *b* sequence is identical to that of a specimen of *H. fuscus* from Safaga on the Red Sea coast of Egypt (Wang et al. 2019a). All Chinese seahorses clustered randomly among sequences of *H. fuscus*, a species name that was considered to be a synonym of *H. kuda* in a recent taxonomic revision of seahorses (Lourie et al. 2016). Bootstrap support for its distinctness was low in both trees, but in the control region tree all *H. fuscus* and *H. casscsio* sequences formed a sister clade to *H. kuda*, and in the cytochrome *b* tree they formed a monophyletic cluster that was nested within a more diverse *H. kuda* cluster. This, and the fact that *H. fuscus* seems to be morphologically distinct from *H. kuda* because of the absence of a coronet (Teske et al. 2005), a trait that it shares with *H. capensis* (Teske et al. 2005) and *H. casscsio* (Zhang et al. 2016), suggests that it is worthy of renewed taxonomic attention. Incomplete lineage sorting between the Arabian Sea and the South China Sea indicates that this species may be represented in the vast, unsampled region separating these two locations. Due to the relatively low information content of the two mitochondrial markers used here, it cannot be ruled out that *H. casscsio* and *H. fuscus* are distinct species that have diverged very recently. However, until this is confirmed with high-resolution genomic data (Teske et al. 2019), the species status of *H. casscsio* should be rejected.

In conclusion, while the generation of mitogenomic data for seahorses can be expected to clarify taxonomic relationships between problematic species, it is important that such resources are compared with as many previously generated sequences as possible prior to publication. In this particular case, the fact that the phylogenetic trees presented in mitogenome announcements typically only include previously published complete mitogenomes (Jooste et al. 2019), to the exclusion of shorter fragments from more closely related species, prevented the discovery of the misidentification of the specimen on which the problematic *H. capensis* mitogenome was based. Moreoever, while the incorporation of genetic data into the species description of *H. casscsio* is commendable, the fact that the new species was only compared with Chinese seahorses, while published data from more closely related species were ignored, likely resulted in nothing more than the description of yet another synonym. This is a problematic trend that we highlighted over a decade ago (Teske et al. 2007). As such, failure to integrate existing molecular data will merely exacerbate the already confusing situation concerning the taxonomy of the genus *Hippocampus*.

**Fig. 1.**
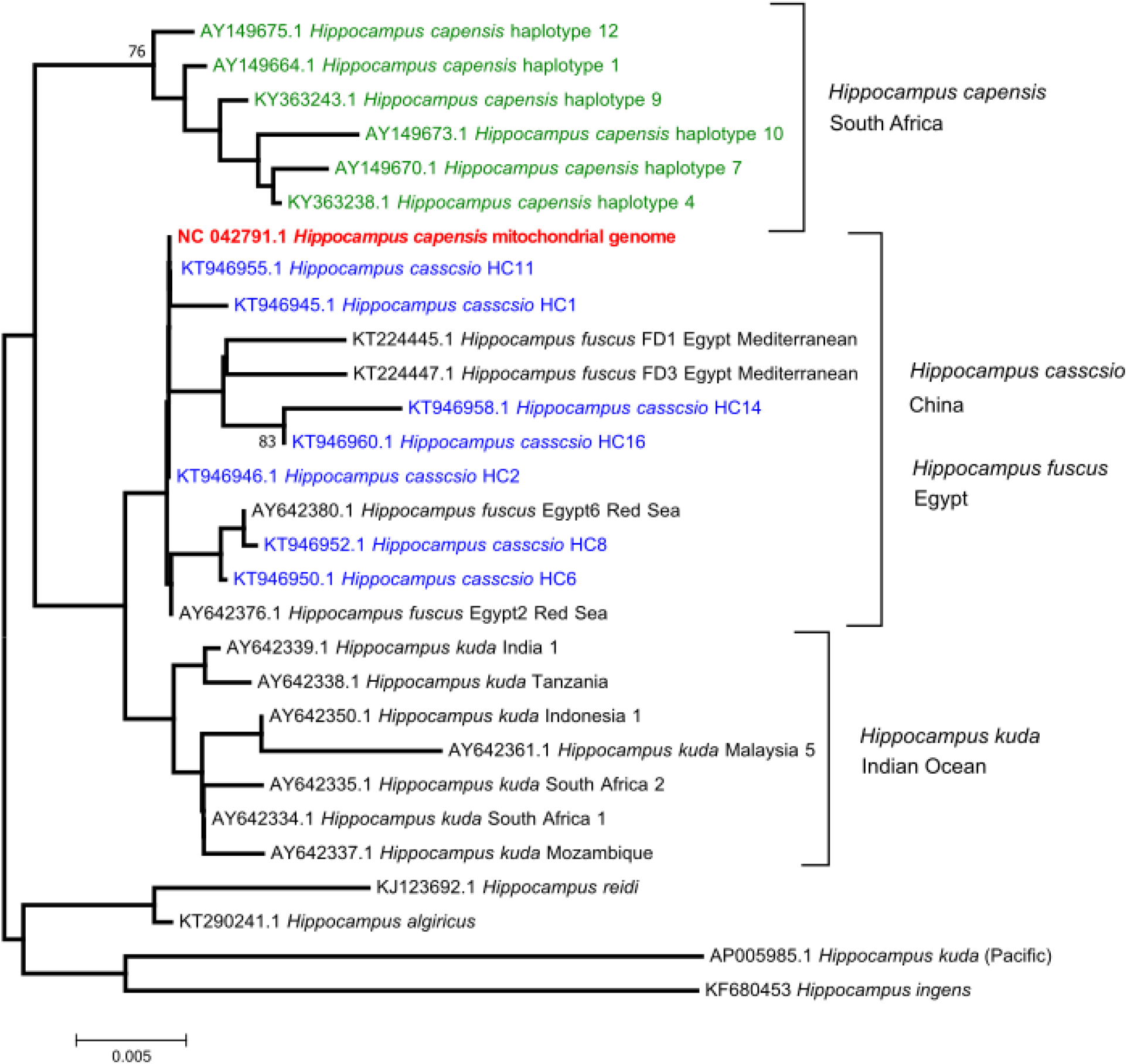
Neighbour-joining tree of 406 bp of control region sequences from *Hippocampus capensis* (South Africa), the Chinese specimen used for the mitogenome reconstruction of *H. capensis* (red), the recently described *H. casscsio* from the South China Sea, and a selection of closely related species. Bootstrap values >75% are shown next to some nodes.

**Fig. 2.**
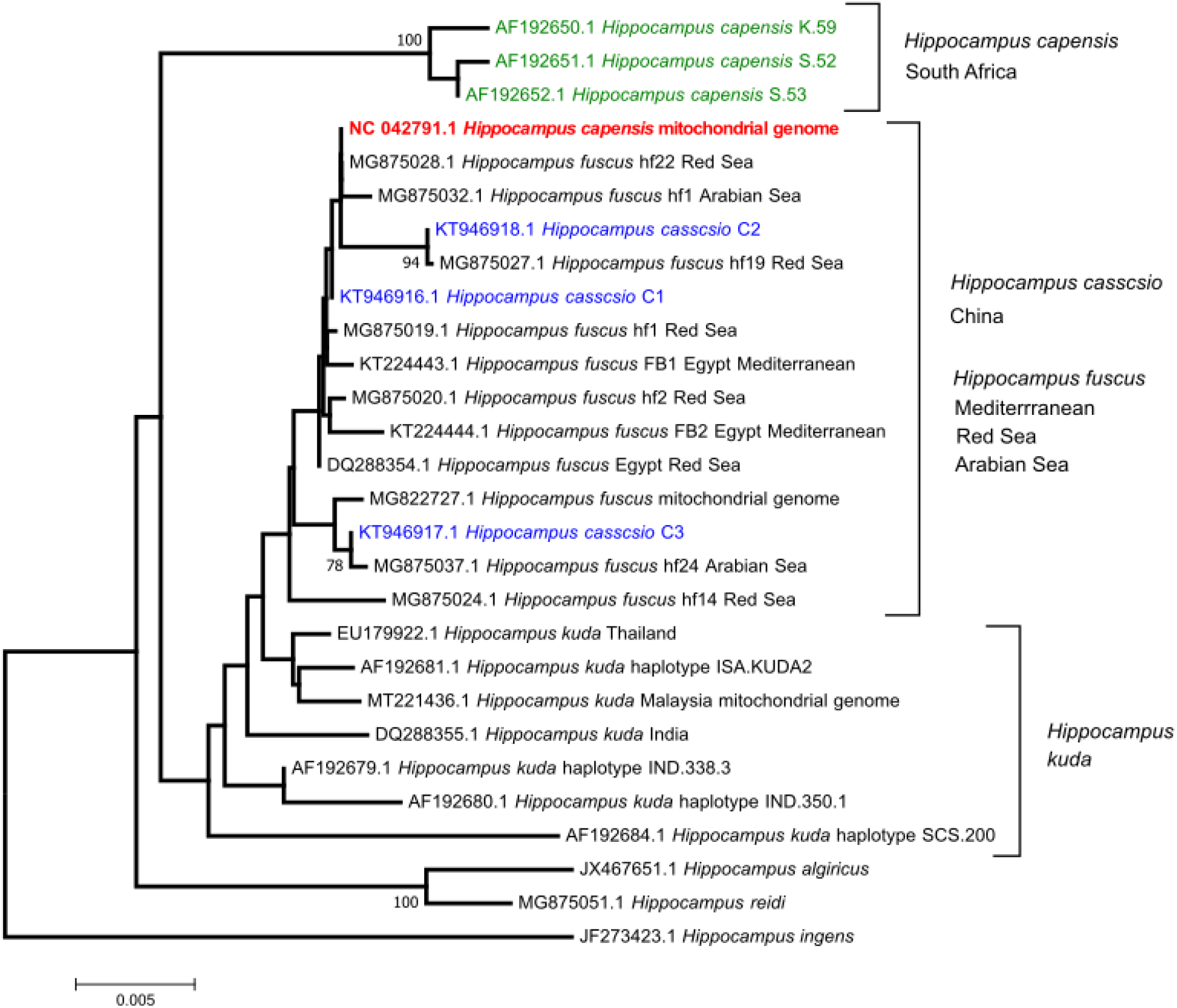
Neighbour-joining tree of 836 bp of control region sequences from *Hippocampus capensis* (South Africa), the Chinese specimen used for the mitogenome reconstruction of *H. capensis* (red), the recently described *H. casscsio* from the South China Sea, and a selection of closely related species. Bootstrap values >75% are shown next to some nodes.

## Acknowledgements

I am grateful to R.A. Sreepada (National Institute of Oceanography, India) for pointing out the problematic mitogenome of *Hippocampus capensis*.

